# EEG-derived Brain Connectivity in Theta/Alpha Frequency Bands Increases During Reading of Individual Words

**DOI:** 10.1101/2025.03.10.642502

**Authors:** Fatemeh Delavari, Zachary Ekves, Roeland Hancock, Gerry T. M. Altmann, Sabato Santaniello

## Abstract

**Objective:** Although extensive insights about the neural mechanisms of reading have been gained via magnetic and electrographic imaging, the temporal evolution of the brain network during sight reading remains unclear. We tested whether the temporal dynamics of the brain functional connectivity involved in sight reading can be tracked using high-density scalp EEG recordings.

**Approach:** Twenty-eight healthy subjects were asked to read words in a rapid serial visual presentation task while recording scalp EEG, and phase locking value was used to estimate the functional connectivity between EEG channels in the theta, alpha, beta, and gamma frequency bands. The resultant networks were then tracked through time.

**Main results:** The network’s graph density gradually increases as the task unfolds, peaks 150-250-ms after the appearance of each word, and returns to resting-state values, while the shortest path length between non-adjacent functional areas decreases as the density increases, thus indicating that a progressive integration between regions can be detected at the scalp level. This pattern was independent of the word’s type or position in the sentence, occurred in the theta/alpha band but not in beta/gamma range, and peaked earlier in the alpha band compared to the theta band (alpha: 184±61.48-ms; theta: 237±65.32-ms, *P*-value *P*<0.01). Nodes in occipital and frontal regions had the highest eigenvector centrality throughout the word’s presentation, and no significant lead-lag relationship between frontal/occipital regions and parietal/temporal regions was found, which indicates a consistent pattern in information flow. In the source space, this pattern was driven by a cluster of nodes linked to sensorimotor processing, memory, and semantic integration, with the most central regions being similar across subjects.

**Significance:** These findings indicate that the brain network connectivity can be tracked via scalp EEG as reading unfolds, and EEG-retrieved networks follow highly repetitive patterns lateralized to frontal/occipital areas during reading.

## 1. INTRODUCTION

Reading is a complex cognitive process that requires efficient network organization and interaction between multiple brain regions [1]. Early work has shown that sight reading a word begins with sequentially activating regions along the ventral processing streams, progresses anteriorly to activate the left inferior temporal and prefrontal areas [2–7], and elicits the formation of functional networks that underlie semantic and contextual integration [8, 9]. More recently, the analysis of resting-state (RS) networks measured via functional magnetic resonance imaging (fMRI) has shown that the connectivity between the left thalamus and right fusiform gyrus positively correlates with sight word reading [10], and proficient reading skills rely on integrating attention-related regions from the left middle temporal area, right hippocampus, and anterior superior temporal sulcus [11, 12]. Vice versa, greater connectivity between the left supramarginal gyrus and the anterior caudate is associated with weaker reading decoding abilities in adults [13], and activation of the right inferior frontal sulcus and middle frontal gyrus has been shown both in adults with left anterior temporal lobe resection [11] and children with dyslexia [14], thus indicating that reading impairment and/or difficulties may lead to compensatory mechanisms that involve hyperactivation of the dorsal striatum and over-integration of the attention networks. Finally, in children with reading difficulties, improvements in reading skills secondary to training have been associated with increased connectivity between visual- and cognitive-control RS networks, decreased segregation between brain regions, and reduction in functionally distinct clusters of activity [15, 16]. Altogether, these studies indicate that the activation of a spatially distributed ensemble of brain regions underpins reading, and alterations to functional connectivity between these regions emerge in the presence of reading difficulties.

To further understand the spatiotemporal pattern according to which these regions are recruited during word-by-word sight reading, magnetoencephalographic and intracranial electrocorticographic studies [2, 6, 17, 18] showed that the neural activity induced by reading peaks in the beta (13-30 Hz) and gamma (30-50 Hz) ranges approximately 170-ms after the stimulus onset (i.e., the time when the word is first displayed) in the ventral occipitotemporal area and then moves to the inferolateral temporal area and the orbitofrontal and left inferior prefrontal areas, where peaks are observed approximately 230-ms and 350-to-400-ms post-stimulus onset, respectively.

At the scalp level, this pattern leads to a negative event-related potential (ERP) in the centroparietal area approximately 400-ms post-stimulus (N400) [19]. Albeit N400 ERPs are not limited to written language, the N400 amplitude has been found to decrease with the ease of semantic processing and integration in word-by-word sight reading [20, 21]. Also, scalp electroencephalography (EEG) has shown lateralized responses to verb generation, with ERPs emerging in the left temporal region [22] and the power spectrum decreasing in the beta and gamma ranges across the left inferior frontal area [23]. Finally, studies [24, 25] have investigated the propensity of EEG-retrieved brain networks to form clusters and exhibit small-world properties during continuous sight reading in children, showing that clusters are formed in the theta (4-8 Hz), alpha (8-13 Hz), and beta ranges, and small-world properties are primarily measurable in the theta and alpha ranges. This trend was partially reverted in dyslexic children, where the propensity to form clusters during reading significantly shifted from the theta/alpha frequency range to the beta band compared to control subjects [24, 25]. Moreover, strong connectivity and reduced neural propagation between the left ventral occipital and temporal cortices were reported in healthy individuals during visual word processing, while dyslexia was reported to disrupt this pathway, increase the propagation time, and elicit a compensatory recruitment of the right posterior brain regions [26, 27]. Altogether, these studies indicate that scalp EEG can detect responses to language tasks and help retrieve the brain networks that are activated during word-by-word sight reading.

It remains unclear, though, how EEG-based brain networks evolve over time and across frequency bands as the word-by-word reading task unfolds. It is also unclear whether the network evolution pattern during sight reading changes depending on the word’s semantics or position within the sentence. Our study addresses both questions using a rapid serial visual presentation task, high-density EEG recordings, and a measure of phase-locking between band-limited EEG time series. We also reconstructed scalp EEG recordings in source space and show that sight reading is associated with the activation of a widespread network involving sensory processing, cognitive, and sensorimotor regions. Differently from previous studies, we show that: (i) a network associated with the reading function can be reconstructed from scalp EEG time series in the range 4-12 Hz; (ii) this network mainly depends on the phase difference between adjacent regions, with this difference showing similar patterns across individuals; and (iii) at a functional level, EEG-based networks undergo significant modulation during reading, with this modulation being largely independent of the word type and syntactic construct.

## 2. MATERIALS AND METHODS

### 2.1. Participants and Reading Experiment

Twenty-eight healthy individuals (male/female: 7/21; age: 18-24 years) who are native English speakers with normal hearing and normal or corrected-to-normal vision were invited to participate in a rapid serial visual presentation reading task. A total of 160 sentences were displayed word-by-word on a computer monitor during the task, with each word being displayed for 300-ms. A 300-ms-long gap between consecutive words was added by displaying a blank screen on the monitor. Participants were asked to read one word at a time until a sentence was completed. Sentences included here were part of an unrelated study that aimed to uncover how the cognitive system tracks and recovers distinct representations of an object following an event described through language, e.g., see [28, 29] for details. Specifically, sentences consisted of two clauses, which detail the “*before*” and “*after*” of an event, respectively, and were organized in four groups (40 sentences per group) depending on the magnitude of the event and how the object is referred to in the sentence, i.e.,

- **Group 1:** The object described in the sentence undergoes a substantial change in state, and its name is repeated in both clauses, e.g., “*The chef will chop the **tomato**. And then he will smell the **tomato**.*”
- **Group 2:** The object described in the sentence undergoes a minimal change in state, and its name is repeated in both clauses, e.g., “*The chef will weigh the **tomato**. And then he will smell the **tomato**.*”
- **Group 3:** The object described in the sentence undergoes a substantial change in state and is indicated by a pronoun in the second clause, e.g., “*The chef will chop the **tomato**. And then he will smell **it**.*”
- **Group 4:** The object described in the sentence undergoes a minimal change in state and is indicated by a pronoun in the second clause, e.g., “*The chef will weigh the **tomato**. And then he will smell **it**.*”

Sentences from group 1-4 were interleaved and presented according to a random order to each participant. The study was carried out in accordance with the Declaration of Helsinki and was approved by the institutional review board at the University of Connecticut, with participants giving written consent prior to engaging in the experiment.

### 2.2. HD-EEG Acquisition and Preprocessing

High-density (HD)-EEG signals were recorded by a 256-channel EGI system, MagStim EGI Inc., Eugene, OR, at a sampling rate of 250 Hz and processed in EEGLAB, ver. 2022.1 [30], and MATLAB, rel. 2021b, The MathWorks, Natick, MA. EEG signals were acquired continuously throughout the presentation of all 160 sentences and digitally filtered offline between 1-Hz and 50-Hz by applying a 256-order high-pass and 60-order low-pass finite impulse response (FIR) filter, respectively (zero-phase filtering). Power line noise at 60 Hz was also removed by using a second-order notch filter, and EEG channels corresponding to electrodes placed on a subject’s face or neck were removed.

Filtered EEG signals were processed in EEGLAB by using the **`Clean_Rawdata`** plug-in, ver. 2.0 [31] to identify bad channels and noisy epochs. A channel was deemed “*bad*” and discarded entirely from our analysis if (i) the ratio between standard deviation (S.D.) and mean of the channel’s EEG time series was greater than 3 and (ii) the Pearson’s correlation coefficient between the EEG time series at that channel and neighbor channels was below 0.7. Conditions (i)-(ii) were determined to balance between sensitivity and specificity in detecting noise. After discarding bad channels, EEG time series from the remaining channels were segmented into 500-ms-long, nonoverlapping epochs, with each epoch starting at the onset of a word’s display, and noisy epochs were identified by using the Artifact Subspace Reconstruction (ASR) method [32], i.e., an epoch was deemed “*noisy*” and discarded from the analysis (i.e., all channels in the epoch were removed) if the S.D. of the multichannel EEG time series in that epoch was greater than twenty-folds the S.D. of the ASR-reconstructed baseline EEG. For each subject, independent component analysis [33] was then performed on the retained EEG epochs to remove muscle, eye, and electrocardiographic components. Putative sources for the independent components were determined by running the **`ICLabel`** plug-in, ver. 1.4 [34] in EEGLAB, and further confirmation was obtained by visually inspecting the scalp topography, ERP image, and power spectrum of each component. Components that were positively confirmed as related to muscle, eye, and/or electrocardiographic activity were removed before reconstructing the EEG time series. Finally, in each epoch, bad EEG channels were reconstructed from the adjacent, clean EEG channels via spherical interpolation using the **`eeg_interp`** function in EEGLAB [35]. All reconstructed EEG signals were then re-referenced (common average referencing) to reject noise across electrodes.

### 2.3. Brain Network Graph Analysis

For each participant, we envisioned the clean EEG channels as nodes in the brain network and hypothesized that the connectivity between nodes evolves over time as the reading task unfolds. Accordingly, we considered the 500-ms-long EEG epochs spanning the display of individual words as separate trials, and we defined the network connectivity as a trial-averaged graph whose topology evolves from the appearance of a word (i.e., *t=*0-ms) on the monitor until the completion of the corresponding cognitive process (i.e., *t=*500-ms; increments: 4-ms), which is set by including 200-ms after the removal of the word from the display. The 100-ms-long gap preceding the display of the following word was removed from our analysis to avoid the contamination with the prediction process of the next word [36].

For any two nodes 1 ≤ *r*, *s* ≤ *N* in the brain network, where *N* is the number of retained EEG channels, we band-pass filtered the EEG time series at the nodes in the assigned frequency band, ℬ, with zero-phase FIR filter of order 150, 100, 50, and 25 for ℬ = *θ* (4-7 Hz), *⍺* (8-12 Hz), *β* (13-30 Hz), and *γ* (30-50 Hz), respectively. Then, we worked on each frequency band ℬ separately and defined the strength of the connection between nodes over time in band ℬ as the phase-locking value (PLV) [37]

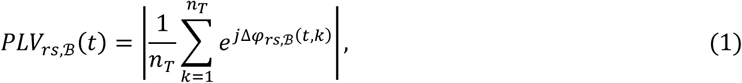

where *n*_*T*_ is the number of trials, Δ*φ*_*rs*,ℬ_(*t*, *k*) = *φ*_*r*,ℬ_(*t*, *k*) − *φ*_*s*,ℬ_(*t*, *k*) is the difference between the instantaneous phases, *φ*, of the band-limited EEG time series at channel *r* and *s*, respectively, at time 0 ≤ *t* ≤ 500-ms in trial *k*, and |·| denotes the absolute value. Since PLV ranges between 0 and 1 and measures the intertrial variability of the phase difference at each time point [37], values for *PLV*_*rs*,ℬ_ (*t*) close to 1 indicate that the phase difference at time *t* is stable across words and reflects synchrony between nodes *r* and *s* at that time of the cognitive process.

We defined one graph per time point *t*, 𝒢_*t*,ℬ_, with the graph only retaining connections that were indicative of synchrony between nodes, i.e., each graph 𝒢_*t*,ℬ_ was defined by an *N*×*N* adjacency matrix whose element in position (*r*,*s*), with 1 ≤ *r*, *s* ≤ *N*, is 1 if *PLV*_*rs*,ℬ_(*t*) passed a statistical test based on bootstrapping and 0 otherwise, see ***Supplementary Text 1***. The topology of the graph 𝒢_*t*,ℬ_ was characterized by measuring the density (𝒟) and average shortest path length (ℒ), which quantify the graph’s compactness and integrity, respectively, and are defined as:

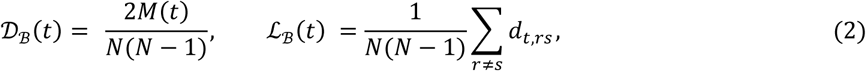

respectively, where *M*(*t*) is the number of connections, i.e., edges, defining 𝒢_*t*,ℬ_, *N* (*N* – 1) is the maximum number of possible edges in the graph, and *d*_*t*,*rs*_ is the length (i.e., number of edges) of the shortest path between nodes *r* and *s* in 𝒢_*t*,ℬ_, for any two nodes 1 ≤ *r*, *s* ≤ *N* with *r* ≠ *s* [38]. Also, the eigenvector centrality (*EVC*_ℬ_) of the nodes [38] was measured by calculating the leading eigenvector of the graph’s adjacency matrix for all *t*, thus tracking the importance of each node over time.

To determine whether the evolution of 𝒟_ℬ_(*t*), ℒ_ℬ_(*t*), and *EVC*_ℬ_(*t*) in time is related to reading, we created one template graph per participant, 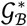, as the graph that retains only the edges that were present in all graphs 𝒢_*t*,ℬ_, with 0 ≤ *t* ≤ 500-ms (i.e., an edge is included in 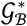 only if it is in 𝒢_*t*,ℬ_ for all *t*), and we calculated the Jaccard similarity coefficient, 𝒮_ℬ_(*t*), between the edge sets of any graph 𝒢_*t*,ℬ_ and the template graph 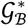. Coefficient 𝒮_ℬ_(*t*) is defined as the size of the intersection between the two edge sets divided by the size of their union [38] and ranges between 0 and 1, with values close to 1 indicating strong similarity between 𝒢_*t*,ℬ_ and 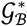, and values close to 0 indicating significant dissimilarity between the graphs, respectively. Accordingly, low values of 𝒮_ℬ_(*t*) indicate that the topological properties of 𝒢_*t*,ℬ_ are substantially different from the template and likely stem from the modulation induced at time *t* by the ongoing reading process.

Finally, for any two nodes 1 ≤ *r*, *s* ≤ *N* and frequency band ℬ, the directed phase lag index [39]

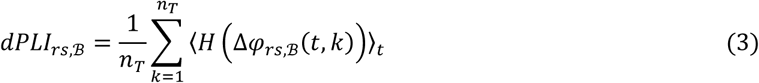

was calculated, where *H*(·) is the Heavyside function, and 〈·〉_*t*_ is the average value over time. Index *dPLI*_*rs*,ℬ_ provides the probability that the instantaneous phase of the band-limited EEG signal at channel *r* is smaller than the instantaneous phase of the signal at channel *s*, and indicates whether channel *r* consistently phase-leads (i.e., 0.5< *dPLI*_*rs*,ℬ_ ≤ 1) or phase-lags (i.e., 0 ≤ *dPLI*_*rs*,ℬ_ < 0.5) compared to channel *s*. The brain network was segmented in four regions (i.e., parietal [*P*], frontal [*F*], temporal [*T*], or occipital [*O*] region), and, for any pair of regions *X*, *Y*, with *X*, *Y=P*, *F*, *T*, or *O*, the average *dPLI*_*rs*,ℬ_ across all pairs of channels (*r*,*s*) with *r* in *X* and *s* in *Y* was calculated to determine the lead/lag relationship between brain regions during reading.

### 2.4. Source Space Analysis

For each participant, the sources (i.e., dipole current density map) of the reconstructed EEG signals were estimated in Brainstorm, ver. 3.4 [40], by using the MNI ICBM 152 atlas, ver. 2009 [41], the cortical surface dipole map (forward model) generated in OpenMEEG [42, 43], and the L_2_-minimum norm estimation method [44]. A total of 210 regions were obtained from the segmentation process, and a mesh grid with 1,082 (scalp), 1,922 (skull), and 1,922 (brain) vertices were used to calculate the dipole currents.

We limited our analysis to a subset of 300 randomly selected epochs (i.e., words) per participant, which correspond to approximately 10% of our dataset, and, for each epoch, we performed the following steps: we first calculated the sources, i.e., dipole currents, at the vertices of the mesh grid (3 sources per vertex) for band-pass filtered EEG signals (band ℬ = *θ*); then, for any pair of regions (*i*, *j*), with 1 ≤ *i*, *j* ≤ 210, we computed the phase-locked value, *PLV*_*ij*_, as the average PLV calculated across all source pairs, with one source located in region *i* and one source located in region *j*; finally, we computed the sample mean (µ) and std. dev. (*σ*) of the values *PLV*_*ij*_ across all regions and epochs and, for each epoch, the connectivity graph between regions was built as a 210×210 binary adjacency matrix whose elements (*i*,*j*) are 1 if *PLV*_*ij*_ > *μ* + *σ* and 0 otherwise. A total of 300 connectivity graphs were built per participant (i.e., one per word), and, for each participant, an edge was deemed significant if it was present in at least half of the graphs.

### 2.5. Statistical Analysis and Part-of-Speech Tagging

To cope with inter-subject fluctuations, we normalized the time series of 𝒟_ℬ_(*t*), ℒ_ℬ_(*t*), and 𝒮_ℬ_(*t*) separately for each participant, i.e., for each participant, the normalized time series *ẑ*(*t*) was derived from the original time series *z*(*t*), with *z* = 𝒟_ℬ_, ℒ_ℬ_, or 𝒮_ℬ_, according to the formula:

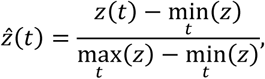

where the minimum and maximum values are calculated separately for each participant. All pair-wise comparisons between graph metrics, i.e., 𝒟̂_ℬ_, ℒ̂_ℬ_, 𝒮̂_ℬ_, or *PLV*_*rs*,ℬ_ (individual edge strengths), in an assigned time window were conducted by using a paired *t*-test (*P*-value *P*<0.05). Tests were conducted for each subject and cumulatively across all subjects. All results are reported as mean ± std. dev. if otherwise indicated.

In case ℬ = *θ* or ℬ = *⍺*, nodes were partitioned between parietal, frontal, temporal, or occipital regions, and the centrality values (i.e., entries in vectors *EVC*_ℬ_(*t*) for all *t*) of nodes in each region were gathered. Similarly, directed phase-lag index values (i.e., *dPLI*_*rs*,ℬ_) were partitioned among all possible pairs of brain regions. Then, differences in importance between regions and inter-region lead/lag relationships during the sight-reading task were tested via ANOVA with Tukey-Kramer *post-hoc* test (*P*-value *P*<0.05) applied to centrality values and directed phase-lag index values, respectively.

In case ℬ = *θ* or ℬ = *⍺*, the time window, *W*, during which graphs 𝒢_*t*,ℬ_ were maximally dissimilar from the template graph 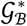 was determined, and differences in graph metrics in window *W* between words was assessed when words are grouped according to type (i.e., *determiner*, *noun*, *auxiliary*, *verb*, *conjunction*, *adverb*, or *pronoun*; tagging conducted in Python by using the **`spaCy`** library, ver. 3.7.5) or role in the sentence (i.e., *subject*, *verb*, or *object* of the first clause, or *subject*, *verb*, or *object* of the second clause). In both cases, differences between word groups were tested via ANOVA with Tukey-Kramer *post-hoc* test (*P*-value *P*<0.05).

## 3. RESULTS

### 3.1. Brain Network Connectivity Evolves through Word-by-Word Sight Reading

A total of 192 EEG electrodes were included in our study, with an average of 29±10 bad channels per participant reconstructed from neighboring clean channels. A total of 2,084±642 trials (i.e., 500-ms-long epochs, with each epoch corresponding to the reading of one word) were retained per participant after removing noisy epochs.

**Fig. 1** shows how the brain network graph in the theta frequency band evolved as the reading of individual words unfolds. The baseline activity, as retrieved by the template graph, 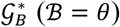, indicates diffused connectivity within the frontal and occipital areas throughout the task (**Fig. 1A**) and limited communication between regions, see **Table 1**, which is consistent with steady activation of the visual cortex during the presentation of visual stimuli and likely represents activity that is not related to reading.

**Figure 1.**
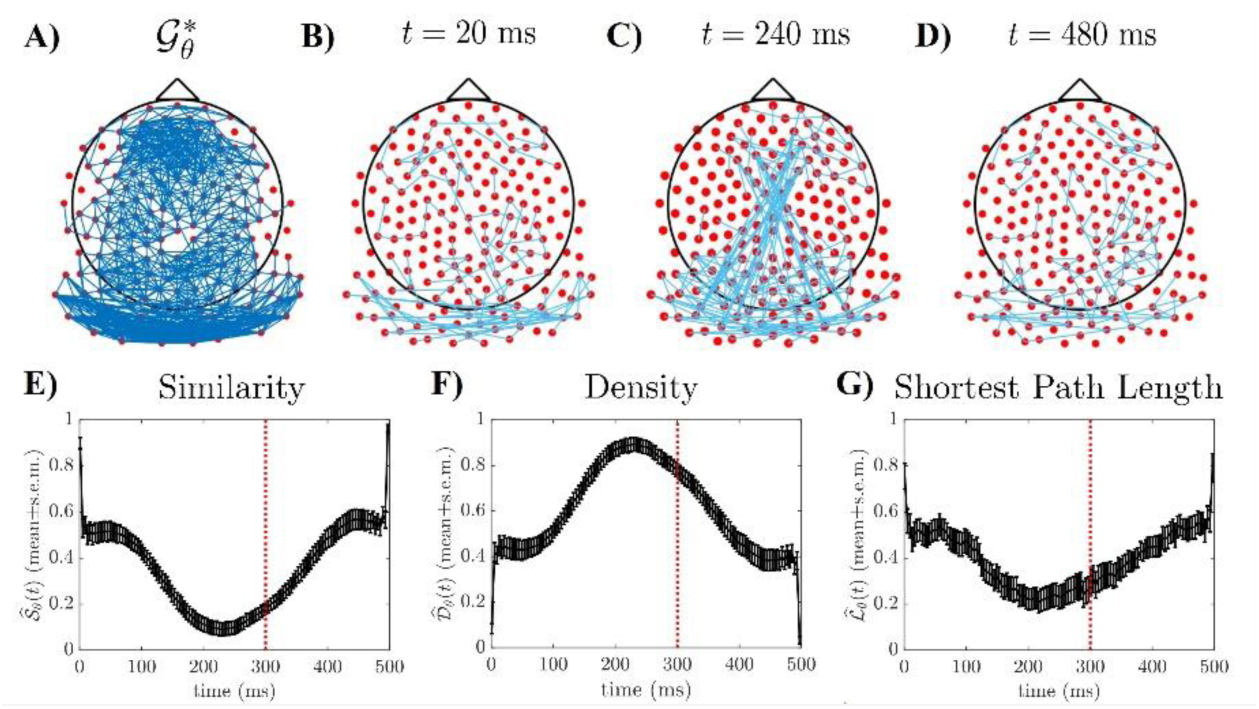
Graph connectivity in the theta frequency band (i.e., ℬ = *θ*) during word-by-word sight reading. A) Template graph (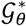) estimated for one participant (participant no. 11) during the presentation of individual words. An edge (*dark blue line*) links any two EEG nodes (*red dots*) if the pairwise PLV between those nodes passed the statistical test indicated in ***Supplementary Text 1*** for the entire duration of the trial, i.e., for all *t*, with 0 ≤ *t* ≤ 500-ms. **B-D)** Difference between 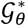 and the graph 𝒢_*t*,*θ*_ estimated at time *t=*20-ms (**B**), 240-ms (**C**), and 480-ms (**D**), respectively, in the same participant. *Light blue* lines in **B-D)** denote edges that selectively emerged in 𝒢_*t*,*θ*_ and are not present in 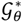. PLV used to determine the edges in **A-D)** were calculated across *n*_*T*_ *=*2,479 trials (i.e., words). **E)** Population-averaged similarity score between graph 𝒢_*t*,*θ*_ and template 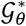 versus time. **F-G)** Population-averaged graph density (**F**) and shortest path length (**G**) of graphs 𝒢_*t*,*θ*_ versus time, respectively. For each time *t* (increments: 4-ms), metrics in **E-G)** are reported as mean ± s.e.m. (*black line* and *vertical black bars*, respectively) across all participants after normalization. *Dashed red* lines in **E-G)** mark time *t=*300-ms, i.e., the removal of each visual stimulus from the screen.

**Table 1.** Density (𝓓_𝓑_) of the template graph 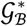 estimated for frequency band 𝓑 = *θ*, *α*, *β*, *γ*. Values are reported for each participant and averaged (“Avg.” row) across participants. Density is calculated separately for each region, i.e., *P*, *F*, *T*, *O*.

| Part.<br>No. | $B = \theta$ | | | | $B = \alpha$ | | | | $B = \beta$ | | | | $B = \gamma$ | | | |
| --- | --- | --- | --- | --- | --- | --- | --- | --- | --- | --- | --- | --- | --- | --- | --- | --- |
| | $F$ | $T$ | $P$ | $O$ | $F$ | $T$ | $P$ | $O$ | $F$ | $T$ | $P$ | $O$ | $F$ | $T$ | $P$ | $O$ |
| 1 | 0.48 | 0.10 | 0.27 | 0.27 | 0.44 | 0.15 | 0.28 | 0.24 | 0.39 | 0.11 | 0.21 | 0.25 | 0.21 | 0.05 | 0.11 | 0.17 |
| 2 | 0.29 | 0.06 | 0.45 | 0.22 | 0.42 | 0.07 | 0.37 | 0.36 | 0.21 | 0.05 | 0.27 | 0.24 | 0.11 | 0.02 | 0.03 | 0.23 |
| 3 | 0.34 | 0.23 | 0.22 | 0.48 | 0.44 | 0.25 | 0.26 | 0.53 | 0.14 | 0.24 | 0.18 | 0.35 | 0.08 | 0.21 | 0.12 | 0.13 |
| 4 | 0.20 | 0.07 | 0.16 | 0.18 | 0.31 | 0.11 | 0.17 | 0.19 | 0.20 | 0.06 | 0.14 | 0.13 | 0.11 | 0.03 | 0.07 | 0.09 |
| 5 | 0.24 | 0.03 | 0.15 | 0.14 | 0.21 | 0.02 | 0.12 | 0.14 | 0.12 | 0.02 | 0.07 | 0.05 | 0.02 | 0.01 | 0.04 | 0.02 |
| 6 | 0.32 | 0.05 | 0.15 | 0.21 | 0.29 | 0.04 | 0.11 | 0.22 | 0.14 | 0.02 | 0.05 | 0.09 | 0.05 | 0.02 | 0.02 | 0.06 |
| 7 | 0.34 | 0.13 | 0.18 | 0.61 | 0.39 | 0.14 | 0.16 | 0.63 | 0.26 | 0.16 | 0.12 | 0.58 | 0.25 | 0.16 | 0.08 | 0.60 |
| 8 | 0.61 | 0.07 | 0.25 | 0.30 | 0.56 | 0.08 | 0.24 | 0.24 | 0.21 | 0.01 | 0.16 | 0.10 | 0.10 | 0.01 | 0.03 | 0.03 |
| 9 | 0.27 | 0.11 | 0.15 | 0.44 | 0.24 | 0.13 | 0.18 | 0.39 | 0.15 | 0.10 | 0.12 | 0.32 | 0.03 | 0.04 | 0.04 | 0.15 |
| 10 | 0.46 | 0.10 | 0.27 | 0.32 | 0.50 | 0.11 | 0.27 | 0.25 | 0.36 | 0.07 | 0.22 | 0.16 | 0.24 | 0.03 | 0.14 | 0.11 |
| 11 | 0.29 | 0.06 | 0.16 | 0.30 | 0.43 | 0.13 | 0.18 | 0.32 | 0.22 | 0.08 | 0.15 | 0.25 | 0.11 | 0.02 | 0.11 | 0.11 |
| 12 | 0.25 | 0.07 | 0.17 | 0.44 | 0.22 | 0.06 | 0.14 | 0.34 | 0.10 | 0.04 | 0.09 | 0.24 | 0.04 | 0.02 | 0.06 | 0.18 |
| 13 | 0.62 | 0.37 | 0.31 | 0.99 | 0.77 | 0.35 | 0.25 | 0.84 | 0.32 | 0.28 | 0.24 | 0.83 | 0.23 | 0.14 | 0.22 | 0.36 |
| 14 | 0.39 | 0.09 | 0.36 | 0.26 | 0.33 | 0.10 | 0.44 | 0.23 | 0.30 | 0.08 | 0.27 | 0.22 | 0.20 | 0.04 | 0.18 | 0.13 |
| 15 | 0.50 | 0.11 | 0.18 | 0.24 | 0.53 | 0.14 | 0.18 | 0.21 | 0.31 | 0.07 | 0.15 | 0.16 | 0.20 | 0.03 | 0.08 | 0.08 |
| 16 | 0.39 | 0.22 | 0.22 | 0.50 | 0.36 | 0.23 | 0.23 | 0.53 | 0.28 | 0.18 | 0.15 | 0.42 | 0.12 | 0.09 | 0.05 | 0.29 |
| 17 | 0.43 | 0.13 | 0.16 | 0.38 | 0.39 | 0.11 | 0.15 | 0.37 | 0.23 | 0.10 | 0.14 | 0.39 | 0.16 | 0.05 | 0.11 | 0.34 |
| 18 | 0.14 | 0.11 | 0.22 | 0.45 | 0.31 | 0.13 | 0.23 | 0.36 | 0.11 | 0.11 | 0.20 | 0.23 | 0.06 | 0.08 | 0.13 | 0.10 |
| 19 | 0.38 | 0.18 | 0.25 | 0.35 | 0.45 | 0.22 | 0.25 | 0.35 | 0.31 | 0.19 | 0.23 | 0.33 | 0.20 | 0.09 | 0.17 | 0.28 |
| 20 | 0.42 | 0.13 | 0.17 | 0.21 | 0.45 | 0.16 | 0.21 | 0.30 | 0.29 | 0.11 | 0.16 | 0.24 | 0.16 | 0.05 | 0.13 | 0.19 |
| 21 | 0.32 | 0.14 | 0.18 | 0.29 | 0.49 | 0.16 | 0.17 | 0.40 | 0.31 | 0.17 | 0.15 | 0.35 | 0.17 | 0.09 | 0.12 | 0.29 |
| 22 | 0.31 | 0.24 | 0.32 | 0.64 | 0.43 | 0.26 | 0.56 | 0.91 | 0.20 | 0.23 | 0.23 | 0.45 | 0.09 | 0.08 | 0.08 | 0.06 |
| 23 | 0.47 | 0.14 | 0.14 | 0.34 | 0.50 | 0.14 | 0.21 | 0.44 | 0.20 | 0.12 | 0.12 | 0.36 | 0.11 | 0.07 | 0.08 | 0.24 |
| 24 | 0.38 | 0.07 | 0.18 | 0.33 | 0.42 | 0.12 | 0.19 | 0.31 | 0.18 | 0.04 | 0.14 | 0.18 | 0.08 | 0.01 | 0.07 | 0.09 |
| 25 | 0.36 | 0.05 | 0.21 | 0.26 | 0.35 | 0.06 | 0.18 | 0.17 | 0.29 | 0.06 | 0.15 | 0.15 | 0.15 | 0.02 | 0.05 | 0.07 |
| 26 | 0.21 | 0.12 | 0.26 | 0.49 | 0.23 | 0.13 | 0.24 | 0.30 | 0.11 | 0.11 | 0.20 | 0.21 | 0.07 | 0.06 | 0.13 | 0.14 |
| 27 | 0.49 | 0.20 | 0.18 | 0.44 | 0.54 | 0.20 | 0.16 | 0.48 | 0.32 | 0.18 | 0.14 | 0.35 | 0.23 | 0.13 | 0.10 | 0.28 |
| 38 | 0.41 | 0.10 | 0.20 | 0.27 | 0.40 | 0.11 | 0.19 | 0.21 | 0.28 | 0.10 | 0.14 | 0.16 | 0.21 | 0.05 | 0.10 | 0.10 |
| Avg. | 0.34 | 0.12 | 0.20 | 0.34 | 0.38 | 0.13 | 0.21 | 0.34 | 0.22 | 0.10 | 0.15 | 0.26 | 0.13 | 0.06 | 0.09 | 0.16 |

As participants engaged in reading the words on display, though, several edges emerged in addition to the edges of the template graph, **Fig. 1B-D**. Although these additional edges were transient, with fluctuating resurgence and disappearance over time, the total number of additional edges grew monotonically from the onset of the word presentation (**Fig. 1B**) to approximately *t=*240-ms (**Fig. 1C**). Then, the number of additional edges monotonically decreased, with the decrement continuing after removing the visual stimulus (i.e., *t=*300-ms) and reaching low steady-state values for *t* in the range 460-480-ms (**Fig. 1D**).

This trend was consistent across participants, even though the total count of edges in the template graph, 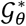, and the number of additional edges brought by graphs 𝒢_*t*,*θ*_ over time varied across participants (average number of edges in 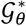: 1,900±472; average number of edges added at time *t=*20-ms, 240-ms, and 480-ms: 159±72, 243±161, and 173±98, respectively; average values calculated across all participants).

To better quantify this trend and closely track the evolution of the network graph over time, we measured the normalized similarity score, 𝒮̂_*θ*_, between graphs 𝒢_*t*,*θ*_ and template graph 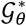 in each participant. **Fig. 1E** shows that, on average, 𝒮̂_*θ*_ remained stable around 𝒮̂_*θ*_= 0.5 in the first 80-ms, then decreased to values close to 0, with the minimum achieved at *t=*237±65-ms (mean± std. dev. across participants), and finally returned to values around 0.5, with the baseline pattern regained well after *t=*300-ms. Correspondingly, the normalized graph density, 𝒟̂_*θ*_, increased as the graphs approached *t=*238-ms (**Fig. 1F**), reduced the average length of the shortest path between nodes, i.e., ℒ̂_*θ*_ (**Fig. 1G**), and then slowly returned to baseline. Also, the trend for 𝒟̂_*θ*_ and 𝒮̂_*θ*_ remained largely unaffected by the selection of the significance level used to screen the phase-locking values and build the graphs, see ***Supplementary Fig. S1A-B***, respectively. Finally, we investigated whether the temporal pattern of 𝒟̂_*θ*_ and 𝒮̂_*θ*_ was specific to sight reading or would rather emerge in response to any visual stimuli on a monitor. Accordingly, we considered scalp EEG time series collected in healthy individuals involved in a picture naming task under similar experimental settings compared to our experiment [45, 46], and we analyzed the EEG-retrieved networks in the theta frequency band, see ***Supplementary Text 2***. Results are summarized in ***Supplementary Fig. S2A-B*** for 𝒟̂_*θ*_ and 𝒮̂_*θ*_, respectively, and indicate that the biphasic shape of the metrics in **Fig. 1E-F** was lost as words organized in structured sentences were replaced with drawings of random objects presented in a random sequence, thus suggesting that the modulatory effect on graph density and similarity was specific to the sight reading task.

Next, we compared the values of 𝒮̂_*θ*_, 𝒟̂_*θ*_, and ℒ̂_*θ*_ in three nonoverlapping widows, i.e., *T*_1_*=*[20, 40] ms, *T*_2_*=*[230, 250] ms, and *T*_3_*=*[460, 480] ms (i.e., 15 graphs per participant altogether), which represent the onset, middle, and end of each epoch, respectively. Windows *T*_1_, *T*_2_, and *T*_3_ were chosen to capture the early fluctuations, the phase of rapid modulation, and the late return to baseline values, respectively, observed in 𝒮̂_*θ*_, 𝒟̂_*θ*_, and ℒ̂_*θ*_, with window *T*_2_ also being the window during which graphs 𝒢_*t*,*θ*_ were maximally dissimilar from the template graph 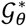. We found significant differences in all three metrics for window *T*_2_ compared to *T*_1_ and *T*_3_ in 25 out of 28 participants and, on average, across all participants (ANOVA with Tukey-Kramer *post-hoc* test, *P*-value *P*<0.01), with no significant difference between windows *T*_1_ and *T*_3_ (*P*>0.05).

### 3.2. Brain Network Connectivity Modulation is Limited to 4-12-Hz Frequency Band

We investigated whether the observed graph connectivity pattern was consistent across frequency bands. **Fig. 2A** shows that the normalized graph density 𝒟̂_ℬ_ increased monotonically up to about *t=*200-ms and then decreased to baseline values in frequency band ℬ = *⍺*, while it presented limited modulation for ℬ = *β*, *γ*. Similarly, the normalized similarity, 𝒮̂_ℬ_, between graphs 𝒢_*t*,ℬ_ and the corresponding template 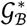 evolved according to a pattern comparable to 𝒮̂_*θ*_ for ℬ = *⍺* but not for ℬ = *β*, *γ*, thus indicating that the observed evolution of graph connectivity during reading was mainly confined to the range 4-12 Hz. This was further supported by the high population-averaged cross-covariance (MATLAB function: **`xcov`**) between 𝒟̂_*θ*_ and 𝒟̂_*⍺*_ compared to the cross-covariance between 𝒟̂_*θ*_ and 𝒟̂_ℬ_, with ℬ = *β*, *γ*, **Fig. 2B**.

**Figure 2.**
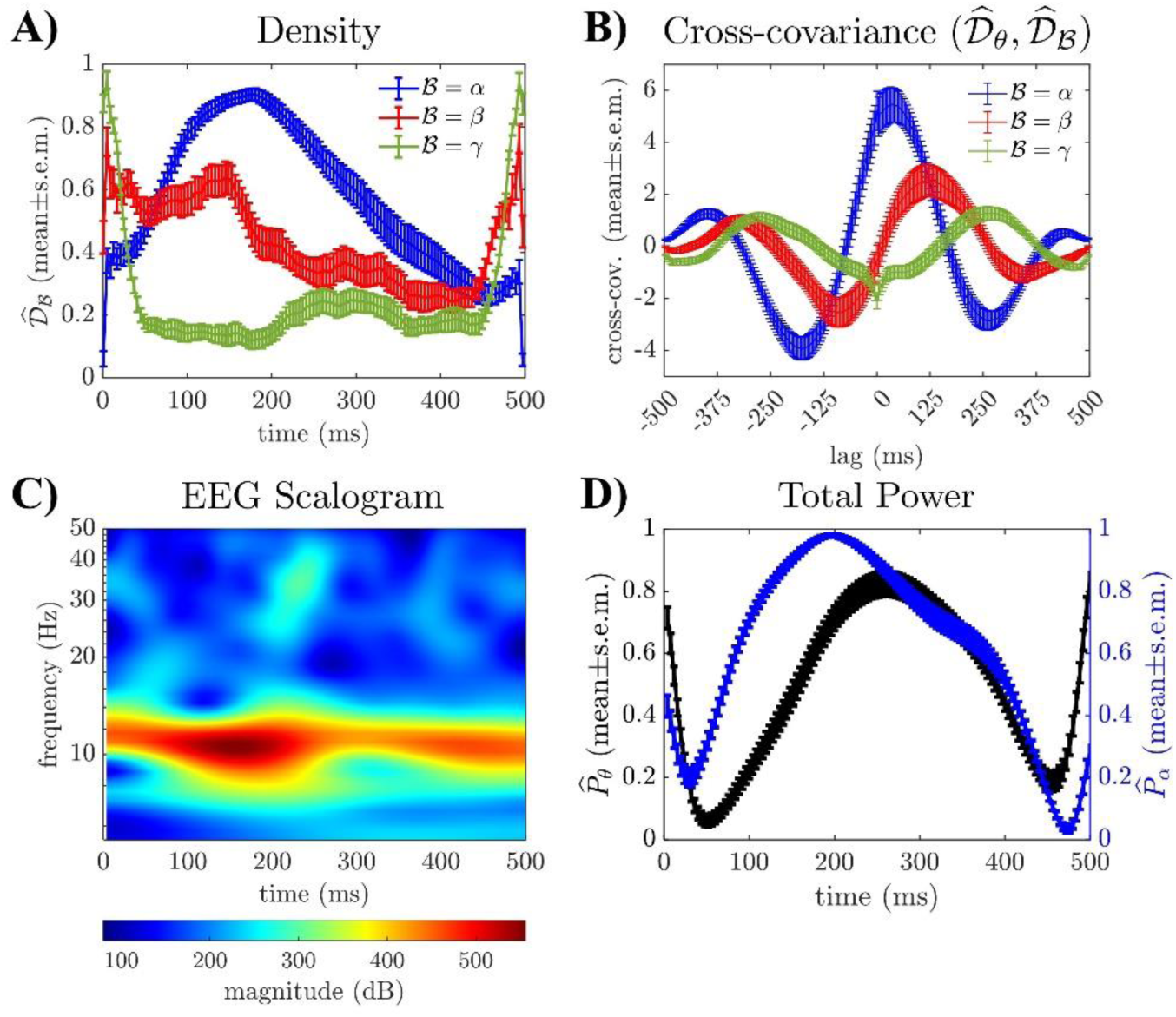
**A)** Population-averaged normalized graph density 𝒟̂_ℬ_ estimated for ℬ = *⍺* (*blue lines*), ℬ = *β* (*red lines*), and ℬ = *γ* (*green lines*), respectively, versus time. **B)** Cross-covariance between the normalized density 𝒟̂_ℬ_ and density 𝒟̂_*θ*_. Cross-covariance is reported as mean ± s.e.m. across participants for each lag. Note that a positive lag indicates that 𝒟̂_ℬ_ precedes 𝒟̂_*θ*_. **C)** Cumulative magnitude scalogram calculated for one participant (i.e., Participant no. 4) during one trial (i.e., one word) across *N=*192 EEG channels. The color scale is shown at the bottom of the plot. **D)** Population-averaged normalized power in the theta (*P̂*_*θ*_, *black lines*) and alpha (*P̂*_*⍺*_, *blue lines*) frequency band versus time. Values are reported as mean (*lines*) ± s.e.m. (*vertical bars*) across all trials and participants.

To further understand the relationship between the graph connectivity in alpha and theta frequency band, we computed the cross-correlation (MATLAB function: **`xcorr`**) between 𝒟_*θ*_ and 𝒟_*⍺*_ and between 𝒮_*θ*_ and 𝒮_*⍺*_ (i.e., values before normalization) in every participant and looked at the lag at which the cross-correlation was maximized. We found that, on average, pairs (𝒟_*θ*_, 𝒟_*⍺*_) and (𝒮_*θ*_, 𝒮_*⍺*_) were highly correlated (max correlation value, normalized to the range 0-1: 0.94 ± 0.05 and 0.88 ± 0.10 for density and similarity, respectively, mean ± std. dev.), with the peak density and peak similarity values consistently occurring in the alpha frequency band (i.e., 𝒟_*⍺*_ and 𝒮_*⍺*_, respectively) prior than the theta frequency band (theta-to-alpha lag: 26.4 ± 32.7-ms and 3.6 ± 14.4-ms for density and similarity, respectively, mean ± std. dev.; max lag: 120-ms and 72-ms for density and similarity, respectively).

Furthermore, we tested whether the graph connectivity pattern observed in the theta and alpha range is reflective of EEG spectrogram. Accordingly, for any trial *k*, we used the wavelet transform (Morlet wavelet function) to compute the scalogram of all 192 EEG channels and then summed the scalogram magnitude across channels. **Fig. 2C** reports the resultant cumulative scalogram (magnitude) for one trial in one participant and shows that the spectrum of the EEG signals remained sustained in the range 4-12 Hz throughout the entire task, with little modulation over time. To further confirm this, we considered one participant at the time and computed the relative power in the theta and alpha frequency band (i.e., *P̂*_*θ*_ and *P̂*_*⍺*_, respectively) as the ratio between the integral of the scalogram magnitude in the theta and alpha band, respectively, and the integral of the magnitude over the range [1, 50] Hz averaged across all trials and channels, see **Fig. 2D**. We found that, across participants, the spectrum changed modestly over time both in the theta and alpha frequency band, with changes mostly responding to the display and removal of the visual stimuli (i.e., in the first 40-60-ms and right after *t=*300-ms). Moreover, moderate-to-high Pearson correlation coefficients were found between the measures of power and graph density in both frequency bands (correlation *P̂*_*θ*_ vs. 𝒟̂_*θ*_: 0.48 ± 0.39; *P̂*_*⍺*_ and 𝒟̂_*⍺*_: 0.75 ± 0.25; mean ± std. dev. across participants), thus suggesting that the evolution of the graph connectivity is only in part related to the spectral modulation at the EEG time series.

Altogether, these results indicate that the brain network becomes denser and more integrated as reading unfolds. The evolution of the brain network, though, is confined to low frequencies (i.e., 4-12 Hz), extends several milliseconds past the removal of the reading input, and is weakly related to the average time-frequency modulation of the EEG time series, thus suggesting that changes in connectivity are likely due to transient synchronization events that localize across different brain regions. The count of these events grows as the task unfolds, with a progression that is faster in the alpha range compared to the theta range.

### 3.3. Contribution of Major Brain Regions to Graph Connectivity During Reading

To further understand whether the change in connectivity during reading caused a shift in information flow through the EEG nodes, we measured the nodes’ importance in the brain network by calculating the eigenvector centrality (*EVC*_ℬ_) for ℬ = *θ* and averaging EVC in four anatomical regions, i.e., frontal, temporal, parietal and occipital.

**Fig. 3A** reports the sample distribution of *EVC*_*θ*_ for the four regions across all epochs and participants and shows that the most important nodes were concentrated in the frontal and occipital regions, thus indicating that (i) the information flow mainly moved along the antero-posterior axis, and (ii) the functional exchange between brain nodes primarily moved through the frontal and occipital regions, with (i)-(ii) being consistent with early electrographic studies [2–7].

**Figure 3.**
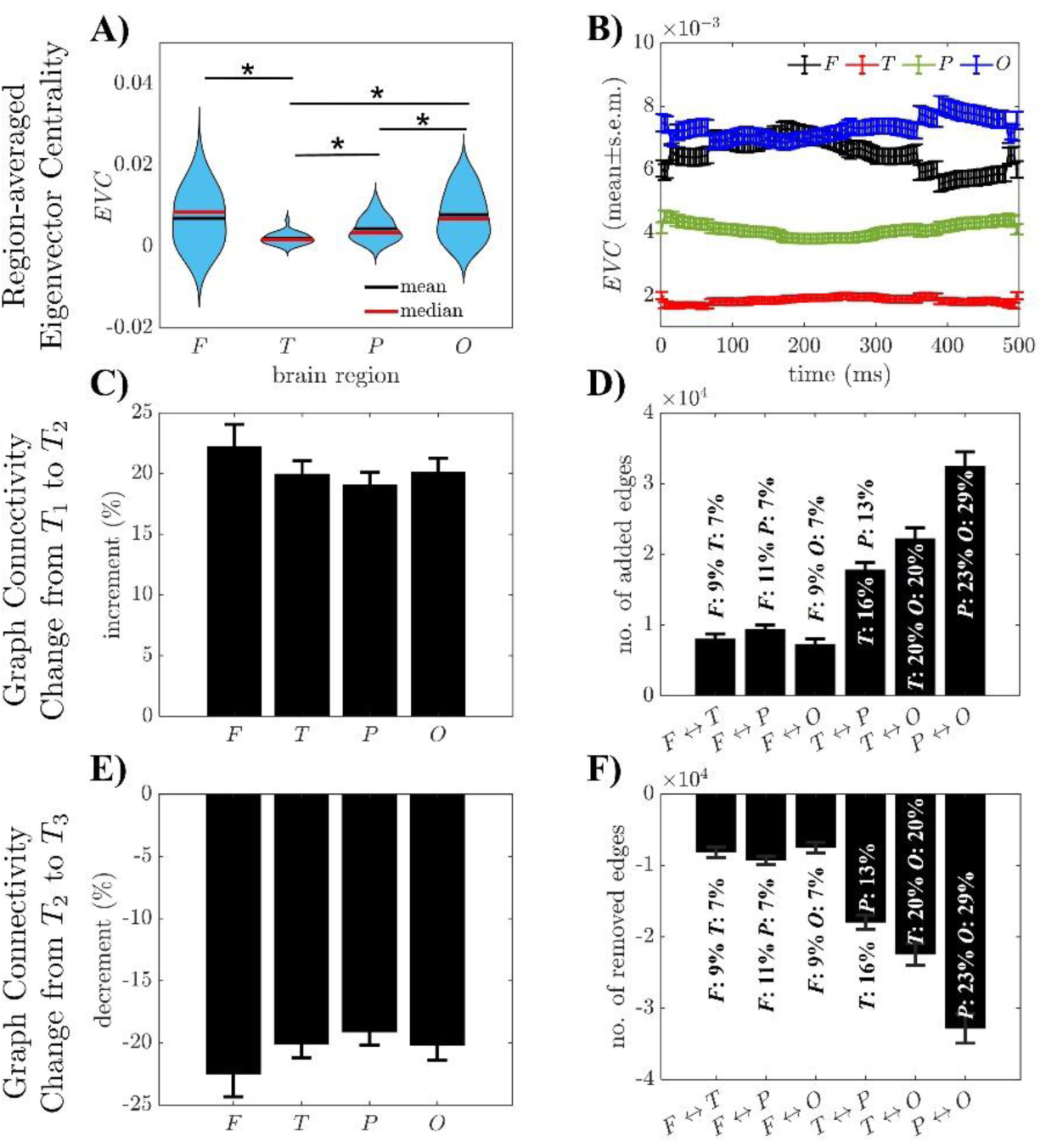
**A)** Violin plots depicting the distribution of eigenvector centrality (*EVC*) values calculated for EEG nodes located in the frontal (*F*), temporal (*T*), parietal (*P*), and occipital (*O*) region of the brain. Values are computed across all participants and epochs. Asterisks denote pair-wise significant difference between regions (*t*-test, *P*-value *P<*0.01). **B)** Population-averaged EVC values in *F* (*black lines*), *T* (*red lines*), *P* (*green lines*), and *O* (*blue lines*) region of the brain vs. time. EVC values are computed at each time *t* (increments: 4-ms) and reported as mean ± s.e.m. (*line* and *vertical bars*, respectively) across all participants. EVC values in **A-B)** are calculated for the frequency band ℬ = *θ*. **C-D)** Increment in edge count of the brain network graphs estimated in window *T*_2_ compared to the graphs estimated in window *T*_1_ for ℬ = *θ* In panel **C)**, increments in edge count in *T*_2_ are reported separately for brain region *F*, *T*, *P*, and *O* as percentage of the edge count calculated for all graphs in *T*_1_. In panel **D)**, increments in edge count in *T*_2_ vs. *T*_1_ are reported separately for edges linking any two brain regions *X* and *Y* (*X*↔*Y*), with *X, Y=F, T, P, O*. For every pair of brain regions *X* and *Y*, the increment in edges linking *X* and *Y* is also reported as percentage of the total edge count involving nodes in region *X* and *Y*, respectively. **E-F)** Decrease in edge count of the brain network graphs estimated in window *T*_3_ compared to the graphs estimated in window *T*_2_ for ℬ = *θ*. In panel **E)**, the decrease in edge count in *T*_3_ is reported separately for brain region *F*, *T*, *P*, and *O* as percentage of the edge count calculated for all graphs in *T*_2_. In panel **F)**, the decreased edge count in *T*_3_ vs. *T*_2_ is reported separately for edges linking any two brain regions *X* and *Y*. For every pair of brain regions *X* and *Y*, the decrease in edges between *X* and *Y* is also reported as percentage of the total edge count involving nodes located in region *X* and *Y*, respectively. Values in **C-F)** are calculated separately for each participant and reported as mean ± s.e.m. across participants.

Furthermore, we measured the average *EVC*_*θ*_ for each region at every time point *t* (**Fig. 3B**) and found that the relative importance of the frontal and occipital regions remained stable throughout the reading task, regardless of the time-varying changes in network density, which indicates that the main hubs of the information flow were largely unaffected by the task-induced temporal evolution. Also, we tested whether the changes in connectivity over time affect the preferred paths of information flow across regions. Specifically, we considered windows *T*_1_, *T*_2_, and *T*_3_ defined in section 3.1 and calculated the number of edges added to each brain region when transitioning from *T*_1_ to *T*_2_ (**Fig. 3C-D**) and removed from each brain region when returning to baseline, i.e., from *T*_2_ to *T*_3_ (**Fig. 3E-F**). We found that, on average, the number of edges added to each region from *T*_1_ to *T*_2_ was close to the number of edges removed from *T*_2_ to *T*_3_, with a slight preference for edges that involve either frontal nodes or occipital nodes (i.e., **Fig. 3C** vs. **Fig. 3E**). On average, the count of edges increased by 20% during the modulation phase of the reading task (i.e., window *T*_2_) in each region, with the increment mostly lost by the time of window *T*_3_. However, when counting edges that were either added (**Fig. 3D**) or removed (**Fig. 3F**) between pairs of brain regions, we found that, on average, the largest fluctuation in edge count occurred between the occipital and parieto-temporal regions (i.e., *O*↔*P*, *O*↔*T*, and *P*↔*T* pairs), with 16-20% temporal nodes, 13-23% parietal nodes, and 20-29% occipital nodes, respectively, first adding and then losing edges throughout the reading process.

Overall, this change in connectivity between occipital, parietal, and temporal areas indicates a significant lateralization of the information flow as the visual stimuli disappears and the reading task is completed. We also investigated the distribution of phase lead/lag between pairs of nodes from adjacent regions via directed phase lag index (*dPLI*) for ℬ = *θ* and found that, on average, no region significantly led nor lagged neighbor regions throughout the reading task (i.e., dPLI values remained close to 0.5 over time with small deviations, see **Table 2**), thus suggesting that, at the network level, the information flow did not follow a preferred orientation as new pathways between occipital and parieto-temporal regions were added.

**Table 2.** Directed phase lag index (𝒅𝑷𝑳***I***) calculated for 𝓑 = *θ* any two regions *X* and *Y* (*X*↔*Y*) with *X*, *Y=F*, *T*, *P*, or *O*. Directed phase lag index values are calculated for each participant and reported as mean ± std. dev. across all participants.

| *Brain Regions* | *$dPLI_\theta$* |
| --- | --- |
| $F \leftrightarrow T$ | $0.502 \pm 0.004$ |
| $F \leftrightarrow P$ | $0.494 \pm 0.010$ |
| $F \leftrightarrow O$ | $0.499 \pm 0.004$ |
| $T \leftrightarrow P$ | $0.496 \pm 0.005$ |
| $T \leftrightarrow O$ | $0.500 \pm 0.006$ |
| $P \leftrightarrow O$ | $0.498 \pm 0.014$ |

To further clarify how functional interactions between distinct brain regions informed the connectivity at the scalp, we projected the EEG channels onto the MNI ICBM 152 atlas and estimated dipole currents across 210 distinct anatomical regions for each participant and epoch. Graphs linking these regions were built for each epoch by using the average PLV between dipole currents in adjacent regions as network edges and only retaining the strongest edges, which resulted in a network topology that varied across epochs and participants, i.e., the same bootstrap method used with EEG time series was applied to sources.

We found that, despite the fluctuations in structure, all graphs identified a similar organization of the information flow across regions. Specifically, for each participant, we measured the importance of the regions in the network by calculating the PageRank centrality [38] of the nodes in every epoch and averaged the PageRank value across epochs. PageRank centrality is a variation of the eigenvector centrality and was preferred to EVC because of the higher number of nodes in the source space compared to EEG channels, which may increase the effect of the nodes’ out-degrees on the node importance [38]. **Fig. 4A** shows the average PageRank centrality of each region in the source space across participants and indicates that a small subset of nodes ranked consistently higher than the remaining nodes (*red dotted line*, which corresponds to the knee-point of the centrality graph), with the list of highest-ranked nodes resulting consistent across all participants and spanning areas mainly located in the left-brain hemisphere, see ***Supplementary Table S1***. Specifically, when placed on the MNI ICBM 152 atlas, the highest-ranked nodes corresponded to the left angular gyrus (rostro-dorsal part) and posterior cingulate cortex (**Fig. 4B**, **(i)-(ii)**), the left supramarginal gyrus (central part) and anterior frontal lobe (**Fig. 4B, (iii)**), and the right fusiform gyrus (dorsolateral part), angular gyrus (caudal and rostroventral parts), lateral prefrontal cortex (**Fig. 4B, (iv)-(v)**), thus demonstrating a consistent recruitment of regions involved in complex language function and cognitive processes.

**Figure 4.**
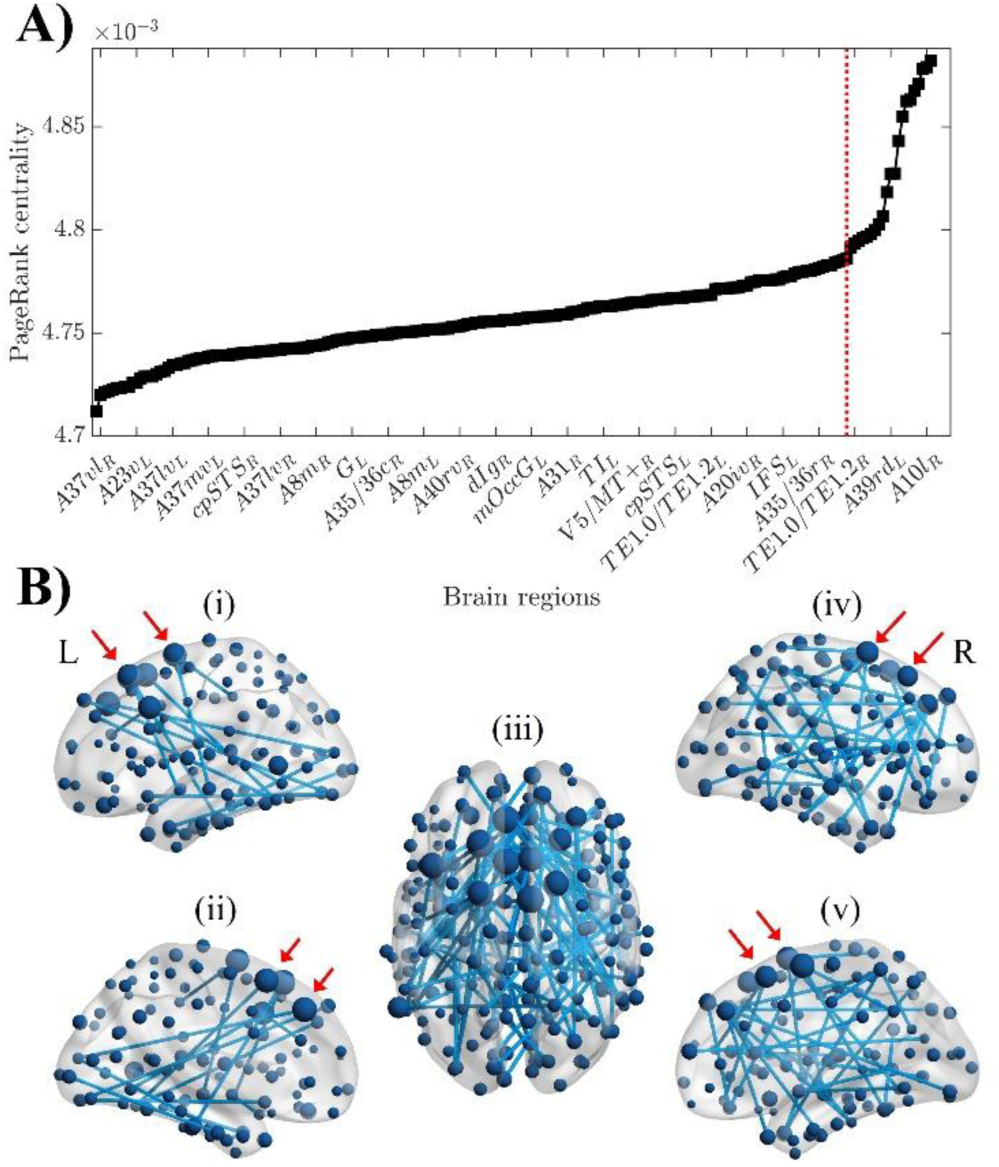
**A)** Population-average PageRank centrality (*black squares*) of the 210 brain regions segmented in the MNI ICBM 152 atlas, with the regions ranked from the least central (left) to the most central (right). A *red dotted line* marks the top 20 regions with the highest PageRank centrality. For each participant, the PageRank centrality of the brain regions is calculated for each trial (i.e., read word) and averaged across trials. The population-average PageRank centrality is calculated as average across participants. **B)** Population-average brain network based on PageRank centrality and overlapped with the MNI ICBM 152 atlas. Overlap generated using the BrainNet Viewer software [47]. Panels (i)-(v) depict sagittal views of the left side (L) of the brain (i-ii), an axial view of the brain (iii), and sagittal views of the right side (R) of the brain (iv-v), respectively. Nodes denote the segmented brain regions, and node sizes are proportional to the PageRank centrality values in **A)**. Edges in panels (i)-(v) depict region-to-region connections that are present in at least 50% of trials in all participants. *Red arrows* indicate some of the top 20 regions listed in **A)**.

Altogether, we found that a widespread network in the theta frequency band can be retrieved from scalp EEG recordings during sight reading, with the structure of this network being lateralized to the left hemisphere, consistent across participants, stable across words, and associated with the selective activation of regions in the source space that are primarily involved in somatosensory representation (e.g., tongue and larynx), semantic integration, and cognitive control.

### 3.4. Impact of Parts-of-Speech on Brain Connectivity during Word-by-Word Sight Reading

Since the evolution of the brain network during sight reading was consistent across words, we wondered whether parts of speech, i.e., the function of words within the sentence, could influence the magnitude of the changes observed in brain connectivity. Hence, we grouped the words in our experiment according to their type and syntactic role in the sentence and, for each word, we counted the number of edges in the brain networks for ℬ = *θ* spanning window *T*_2_, i.e., the window during which the maximum modulation in brain connectivity was observed. Then, we averaged the edge count for each group of words.

**Table 3** reports the results for words grouped according to type (i.e., words in sentences of group 1-4 could be a “*determiner*”, “*noun*”, “*auxiliary*”, “*verb*”, “*conjunction*”, “*adverb*”, or “*pronoun*”) and shows that the average edge count was significantly higher for conjunctions, adverbs, and pronouns compared to nouns and verbs (ANOVA with Tukey-Kramer *post-hoc* test, *P*-value *P*<5×10^-4^). This trend was consistent in 26 out of 28 participants, and nouns frequently presented the lowest number of connections among the seven types. Moreover, in 21 out of 28 participants, the part of speech with the highest number of edges was one among the conjunction, adverb, and pronoun, thus indicating a significant shift toward tokens that (i) have a meaning that depends on the context within which tokens are used, and (ii) may require additional memory recollection and cognitive workload to be processed.

**Table 3.** Average edge count for graphs spanning window *T*2 and calculated for 𝓑 = *θ*. The edge count is reported separately for every type of word. For each word type and participant, the edge count is averaged across all graphs in window *T*2 (i.e., 15 graphs altogether) and available words for that type. Values are reported as X (Y), where X is the average edge count, and Y is the rank of the word type between “*Determiner*”, “*Noun*”, “*Auxiliary*”, “*Adverb*”, “*Conjunction*”, “*Pronoun*”, and “*Verb*”.

| Participant | Type of word |  |  |  |  |  |  |
| --- | --- | --- | --- | --- | --- | --- | --- |
|  | *Determiner* | *Noun* | *Auxiliary* | *Verb* | *Conjunction* | *Adverb* | *Pronoun* |
| 1 | 813 (4) | 881 (3) | 384 (7) | 451 (6) | 642 (5) | 979 (2) | 1520 (1) |
| 2 | 407 (5) | 366 (6) | 360 (7) | 593 (2) | 547 (3) | 532 (4) | 604 (1) |
| 3 | 1498 (2) | 1431 (3) | 1258 (5) | 1875 (1) | 1400 (4) | 1129 (6) | 1008 (7) |
| 4 | 360 (6) | 586 (3) | 326 (7) | 443 (5) | 526 (4) | 855 (1) | 685 (2) |
| 5 | 382 (3) | 274 (7) | 381 (4) | 286 (6) | 636 (1) | 550 (2) | 355 (5) |
| 6 | 384 (4) | 325 (6) | 293 (7) | 327 (5) | 494 (2) | 659 (1) | 443 (3) |
| 7 | 746 (5) | 339 (7) | 1341 (2) | 741 (6) | 754 (4) | 2380 (1) | 932 (3) |
| 8 | 747 (5) | 1066 (3) | 587 (6) | 952 (4) | 441 (7) | 1461 (2) | 1579 (1) |
| 9 | 346 (6) | 334 (7) | 538 (2) | 438 (4) | 770 (1) | 443 (3) | 428 (5) |
| 10 | 865 (5) | 1023 (3) | 560 (7) | 1141 (1) | 868 (4) | 812 (6) | 1118 (2) |
| 11 | 618 (2) | 450 (3) | 447 (4) | 377 (5) | 837 (1) | 364 (7) | 372 (6) |
| 12 | 933 (6) | 2678 (2) | 1193 (5) | 697 (7) | 4230 (1) | 1645 (4) | 2140 (3) |
| 13 | 1339 (4) | 1364 (3) | 910 (7) | 1741 (1) | 1277 (5) | 1522 (2) | 974 (6) |
| 14 | 467 (7) | 664 (5) | 488 (6) | 813 (2) | 903 (1) | 715 (3) | 667 (4) |
| 15 | 733 (4) | 511 (6) | 584 (5) | 359 (7) | 780 (2) | 878 (1) | 765 (3) |
| 16 | 839 (5) | 699 (7) | 786 (6) | 888 (4) | 1358 (1) | 1212 (2) | 1158 (3) |
| 17 | 1166 (3) | 546 (5) | 1490 (1) | 499 (6) | 857 (4) | 296 (7) | 1193 (2) |
| 18 | 543 (3) | 630 (1) | 449 (6) | 509 (4) | 603 (2) | 407 (7) | 472 (5) |
| 19 | 603 (5) | 502 (7) | 622 (4) | 630 (3) | 710 (1) | 642 (2) | 580 (6) |
| 20 | 375 (4) | 306 (7) | 356 (5) | 356 (5) | 438 (3) | 538 (2) | 772 (1) |
| 21 | 591 (3) | 284 (7) | 448 (6) | 451 (5) | 523 (4) | 697 (1) | 625 (2) |
| 22 | 754 (6) | 594 (7) | 919 (4) | 1050 (3) | 1288 (1) | 909 (5) | 1127 (2) |
| 23 | 696 (5) | 414 (7) | 723 (3) | 888 (2) | 699 (4) | 490 (6) | 1603 (1) |
| 24 | 502 (6) | 556 (3) | 524 (5) | 448 (7) | 946 (1) | 528 (4) | 608 (2) |
| 25 | 1218 (4) | 909 (6) | 1291 (2) | 932 (5) | 1236 (3) | 2097 (1) | 667 (7) |
| 26 | 524 (5) | 468 (6) | 335 (7) | 533 (4) | 1044 (3) | 1330 (1) | 1136 (2) |
| 27 | 453 (3) | 394 (7) | 400 (6) | 430 (5) | 437 (4) | 481 (2) | 515 (1) |
| 28 | 380 (6) | 345 (7) | 431 (4) | 430 (5) | 592 (2) | 1129 (1) | 432 (3) |

To further clarify this, we considered the portion of sentences in group 1-4 that orderly includes subject, verb, and object of both clauses and the two bridge tokens between clauses, which were always conjunctions (i.e., “*and*” and “*then*”). Then, we measured the average edge count of the graphs spanning window *T*_2_ for these words according to the order of presentation to the participants, i.e., by separating bridge tokens, first clause, and secondary clause. We found that the edge count associated with bridge tokens was consistently higher compared to the remaining parts of the sentence, especially words forming the first clause, **Fig. 5A**. Also, the third highest edge count in the sentence was consistently associated with the subject of the secondary clause, which was a pronoun by design in all sentences, thus indicating that the higher graph density observed for conjunctions, adverbs, and pronouns was likely related to the additional workload that is required to recall the unfolding of events in the first clause while reading the secondary clause.

**Figure 5.**
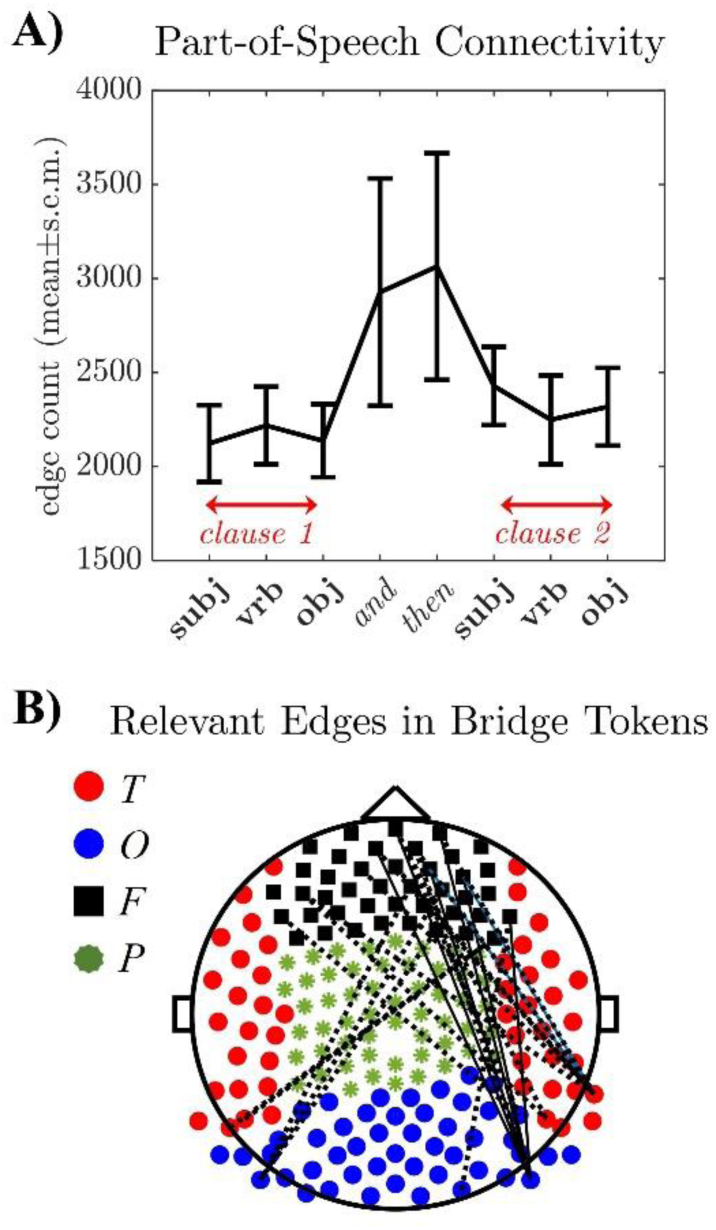
**A)** Average edge count of the graphs calculated in window *T*_2_ for ℬ = *θ* and words that are presented consecutively in a sentence. Sentences in group 1-4 are organized to provide subject, verb, and object of the first clause, two bridge tokens to the secondary clause (i.e., “*and*” and “*then*”), and subject, verb, and object of the secondary clause. Values are averaged across graphs and words per participant and reported as mean ± s.e.m. across participants. **B)** Edges whose average phase-locking value across graphs spanning window *T*_2_ and ℬ = *θ* is significantly higher for bridge tokens (i.e., “*and*” or “*then*”) compared to subject of the first clause. Edges are indicated by *dashed black lines*. Nodes in the frontal (*F*), parietal (*P*), temporal (*T*), and occipital (*O*) region are denoted by *black squares*, *green ribbons*, *red circles*, and *blue circles*, respectively.

This was further confirmed when analyzing the magnitude of the PLV between EEG nodes. Specifically, for each pair of EEG nodes, we compared the phase-locking values calculated for graphs spanning window *T*_2_ (frequency band ℬ = *θ*) while reading the bridge token “*and*” versus the subject of the first clause (paired *t*-test, *P*-value *P*<0.01); then we repeated the same analysis for the bridge token “*then*” versus the subject of the first clause. **Fig. 5B** reports the pairs of EEG nodes that, across all participants, showed phase-locking values that were significantly higher during “*and*” or “*then*” compared to the subject of the first clause. The edges whose strength grew the most during the presentation of bridge tokens (*dashed lines*) were found to connect the parieto-frontal region to the temporal and occipital regions. The spatial arrangement of these edges presented significant lateralization toward regions that are linked with memory, language processing, and cognition (i.e., see sources in **Fig. 4B** corresponding to these nodes).

Altogether, these findings indicate that, albeit different parts of speech have a modest impact on the EEG-based networks, the magnitude of the brain connectivity at the scalp level varies nonlinearly as the sentence unfolds, and the strength of the connections selectively grows at the junction between primary and secondary clauses, i.e., at the point where memory recollection and/or additional cognitive workload is expected to comprehend and track the sentence’s unfolding.

## 4. DISCUSSION

This study provides a characterization of the brain network during word-by-word sight reading and demonstrates that the network’s evolution can be tracked noninvasively through time as reading unfolds by using scalp EEG and a measure of pairwise phase-locking. We also show that intrinsic phase lags between EEG nodes are modulated by sight reading, with the modulation being mainly constrained to the theta/alpha range and largely unaffected by the word’s features. Finally, we show that the magnitude of the modulation of the network increases at the junction between consecutive clauses in complex sentences.

These results are consistent with early studies involving time-frequency analysis of scalp EEG recordings in adults, which showed that neural activation in the theta range is associated with semantic processing. Specifically, increments in EEG theta power have been reported during reading words in isolation, word pairs, or sentences and have been related to the process of retrieving semantic meaning from memory [48–50]. Also, larger increments in theta power during reading have been associated with more difficult retrieval [51, 52] and semantic unification [53]. Finally, it has been shown that increments in EEG theta power while reading are broad across both hemispheres in children (age: 8-9 years) and later lateralize to the left central-posterior areas in adolescents (i.e., age: 12-15 years) [54–57], which suggest that the cognitive system supporting semantic retrieval might not be fully developed by at least age 12 [57].

Our results complement these findings by showing that, in young adults, strong phase-locking between parietal, occipital, and frontal areas preferentially occurs during semantic unification (i.e., at the bridge between clauses), and a diffused front-parietal connectivity emerges during word-by-word reading. This suggests that semantic retrieval and unification evoke an extensive coordination (i.e., phase-locking) between several regions, with this transient coordination not necessarily reflecting a generalized exaggeration of theta oscillations. This is further confirmed by the source space localization of our EEG time series, which indicates significant centrality for regions involved in somatosensory representation and cognitive control. Hence, it is plausible that, while lateralized theta power increments are a direct consequence of increased neural activation in left posterior and prefrontal cortices, the increased connectivity at the scalp reflects stronger inputs from these cortices to the surrounding regions. This is further supported by recent studies investigating resting state networks retrieved from scalp EEG recordings, which have reported that hubs in the left temporal and parietal cortices in the theta and alpha frequency bands can be associated with reading proficiency [58, 59], and lower connectivity between these hubs and the rest of the resting state network emerge in dyslexic subjects, both children [60] and adults [61], whereas lower coherence between frontal and parietal networks in the alpha and beta bands was reported in children with reading disability compared to healthy controls [62].

Interestingly, we also show similar evolutions for phase-locking-based networks retrieved in the theta and alpha frequency band, despite the alpha rhythm being largely associated with attention, e.g., [63] for a review. In RSVP word recognition tasks, event-related desynchronization in the alpha band has been reported [64], with the magnitude of the desynchronization (i.e., amplitude of the N1 component of the event-related potential 200-280ms post-stimulus) depending on the lexical processing load of the word being read as well as the order according to which words are presented [65]. Also, word recognition-related potentials have been reported mainly in the occipital regions [66], and the shape of these potentials has been shown to include three consecutive stages, i.e., an initial short-lasting burst in post-stimulus alpha power, a transient decrease in induced alpha power, and a final sustained increase [66], where this pattern is believed to reflect the integration process of foveally and parafoveally information acquired during fixation of the word on the screen [67]. Our results further confirm that phase-locking in the alpha range heavily involves nodes covering the occipital region, and the high rate of edges between occipital nodes in our template graphs indicates that the occipital region remains highly active throughout the RSVP task as fixation is maintained. Also, the presence of a peak in connectivity earlier compared to the theta frequency band suggests that the synchronization might be related to the initial short-lasting burst in post-stimulus alpha power shown in the event-related potentials [66], thus indicating that the temporal evolution of the brain connectivity in the alpha frequency band may be affected by the oculomotor behavior.

In addition to this, however, we also found that the phase-locking connectivity is lateralized toward the left frontoparietal areas in the alpha frequency band, with the widespread connectivity lasting beyond the display of the words on the screen. These findings suggest that, while power may increase occipitally, the phase-locking between alpha oscillations is not spatially confined to the occipital area and likely reflects a large-scale coordination between regions on different time scales during semantic retrieval and unification, presumably mediated by synchronous waves propagating along the poster-anterior direction [68].

## 5. CONCLUSIONS

We tracked the temporal evolution of the brain connectivity during word-by-word reading and showed that the brain network in the theta and alpha band follows a bimodal pattern, i.e., it evolves from a clustered, small-world architecture to a densely connected topology, where nodes in the left prefrontal and occipital regions maintain the highest centrality, before returning to the resting-state architecture. We also found that the modulation of the network connectivity remains largely unaffected by the word’s features but increases at the junction between consecutive clauses, thus indicating involvement of these networks with semantic retrieval and unification. Altogether, our results suggest the emergence of transient, band-limited synchronization throughout the brain network as the semantic retrieval unfolds, with the nodes in the left frontal and occipital regions mainly steering the information flow by serving as coordinating hubs in the network.

## AUTHOR CONTRIBUTIONS

F.D.: Conceptualization, Data Analysis, Methodology, Software, Visualization, Writing (Original Draft Preparation).

Z.E.: Experimental Design and Implementation, Data Acquisition.

R.H.: Conceptualization, Methodology, Resources, Supervision, Writing (Review & Editing).

G.T.M.A.: Conceptualization, Methodology, Resources, Supervision, Writing (Review & Editing).

S.S.: Conceptualization, Funding Acquisition, Methodology, Project Administration, Resources, Supervision, Visualization, Writing (Review & Editing).

## ACKNOWLEDGEMENTS

This work was partly supported by the US NSF (National Science Foundation) CAREER Award 1845348 to S.S. The funding agencies had no role in study design, data collection and analysis, decision to publish, or preparation of the manuscript.

## DATA AVAILABILITY STATEMENT

The code developed as part of this study will be made available on GitHub upon acceptance.

## SUPPLEMENTARY INFORMATION

### Text 1: Procedure to Test the Statistical Significance of Phase Locking Values

For any frequency band ℬ, time 0 ≤ *t* ≤ 500-ms, and pair of EEG channels 1 ≤ *r*, *s* ≤ *N*, the significance of the phase-locking value *PLV*_*rs*,ℬ_ (*t*) was determined using a bootstrap method. Specifically, we hypothesized that a bias applies to all phase-locking values because of the application of visual stimuli (i.e., words displayed on the monitor), and that such bias is likely similar across participants and 500-ms-long epochs. Accordingly, the bootstrap method consisted of randomly selecting 10 pairs of EEG channels (“*null pairs*”), with each pair including EEG time series from two randomly selected participants, i.e., each pair gathered recordings from different participants. For each null pair, EEG time series were pre-processed as described in the *Main Text*, divided into 500-ms-long epochs (i.e., each epoch started at the onset of a word’s display and lasted 200-ms after the removal of the word from the screen), and epochs were treated as independent trials. Finally, phase-locking values were calculated for every null pair, trial, and time *t* and then gathered to form the null distribution. One null distribution per frequency band ℬ and time *t* was accordingly created. Any phase-locking value *PLV*_*rs*,ℬ_(*t*) was deemed significant if the hypothesis that *PLV*_*rs*,ℬ_ (*t*) is generated by the null distribution was rejected with 99% confidence (i.e., *P*-value *P*<0.01).

**Figure S1.**
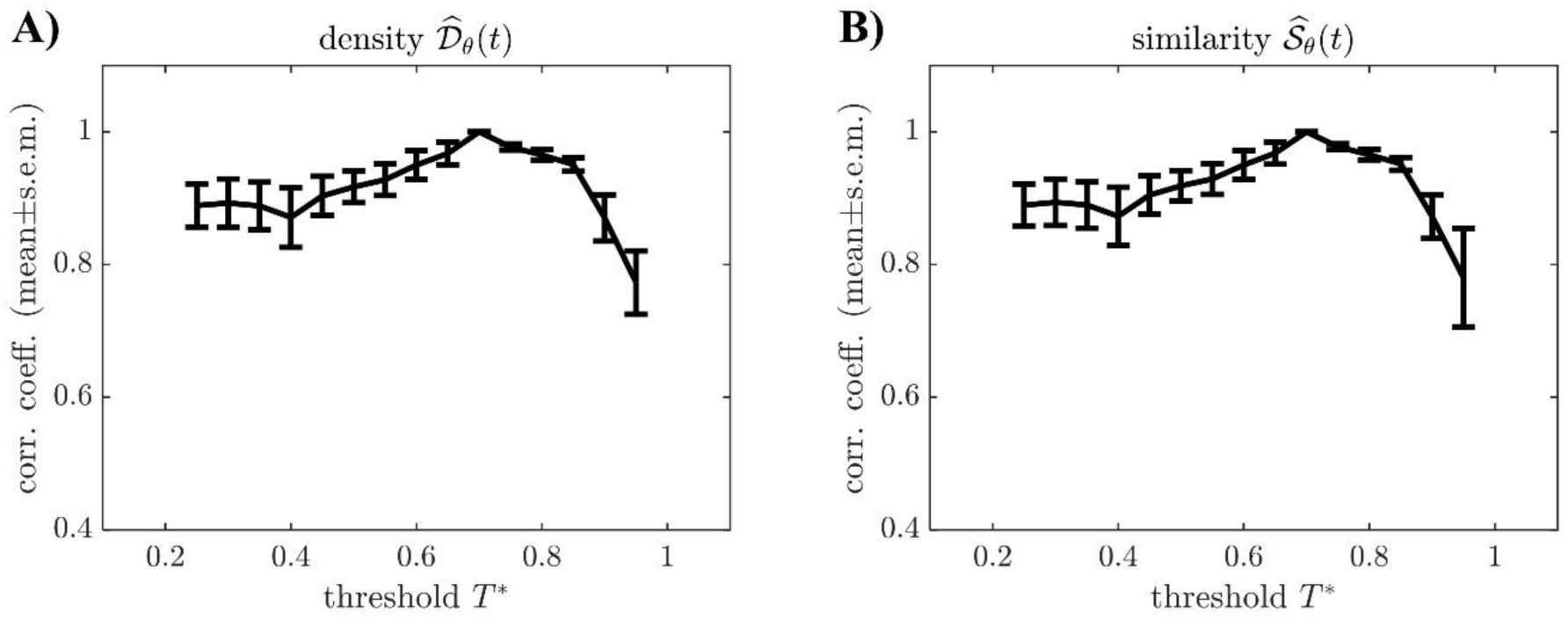
Pearson’s cross-correlation coefficient calculated for the normalized values of density (𝒟̂_*θ*_, **A**) and similarity (𝒮̂_*θ*_, **B**) in the theta frequency band defined in the *Main Text* when the significance of the phase-locking values was determined by using the bootstrap method and when phase-locking values were deemed significant if higher than a threshold *T*^∗^, with *T*^∗^varied from 0.25 and 0.95. Cross-correlation coefficients were computed separately for every participant and reported here as mean ± s.e.m. across participants.

### Text 2: Procedure to Test whether Graph Connectivity Patterns are Specific to Sight Reading

EEG Dataset 1 in [45, 46] consists of multichannel EEG recordings collected from 23 healthy individuals (11 males and 12 females, age: 19-40 years) involved in a picture naming task. Briefly, every participant was seated approximately 1-m in front of a 17-inch screen in a dimly lit room and asked to look at pictures of black drawings on a white background displayed on the screen. For each picture, participants were asked to name the depicted item aloud. A total of 148 drawings, representing examples of two separate categories, i.e., “animals” and “tools” (74 drawings per category altogether), were displayed per participant, arranged in two sessions of approximately 8 min each (74 pictures per session). Participants were asked to name pictures at a normal speed, which resulted in having each picture displayed on the screen for at least one second, with breaks between pictures, and pictures were randomly selected for each participant from a database of 400 images standardized for the French language. Finally, the picture selection process was designed to ensure that every set of 148 images includes items whose name is a short word of 3-to-5 letters (74 images, i.e., 37 animals and 37 tools) and items whose name is a long word of 7-to-10 letters (74 images, i.e., 37 animals and 37 tools).

**Figure S2.**
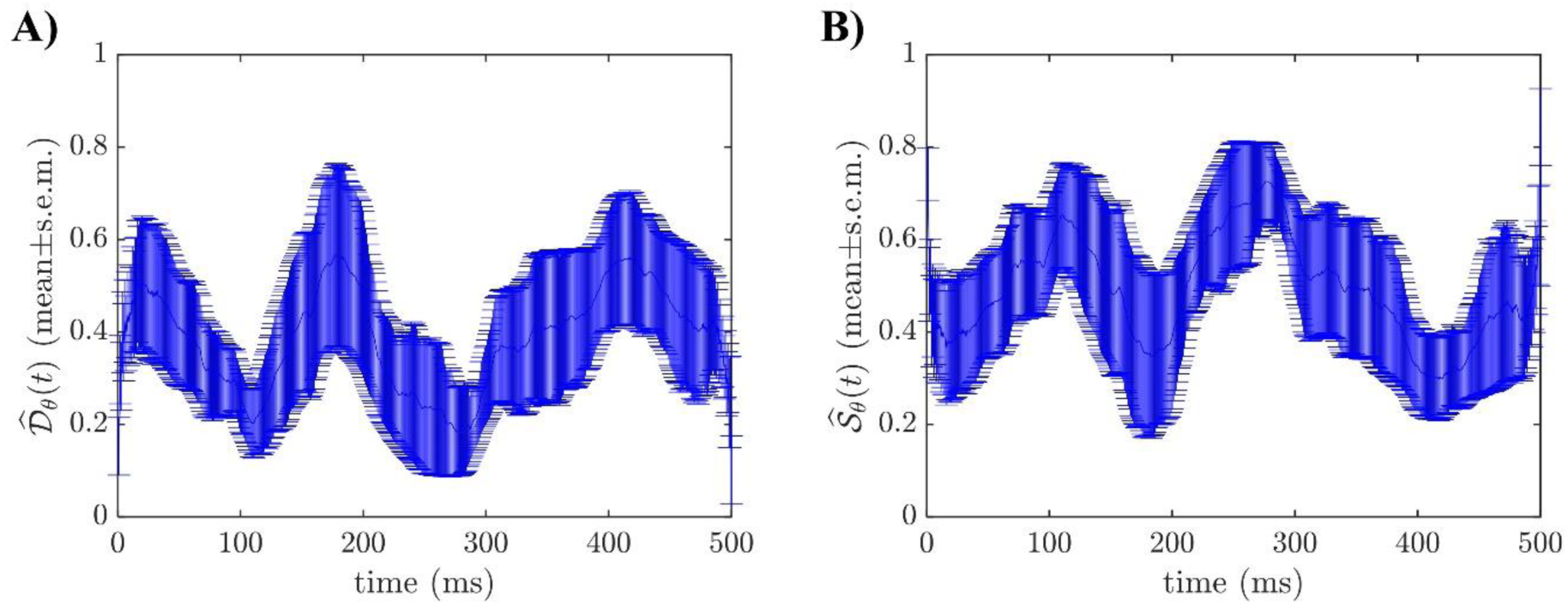
**A)** Population-averaged similarity score between graphs 𝒢_*t*,*θ*_ and template 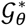 calculated for EEG Dataset 1 versus time. **B)** Population-averaged graph density of 𝒢_*t*,*θ*_ calculated for EEG Dataset 1 versus time. For each time *t* (increments: 4-ms), metrics in **A-B)** are reported as mean ± s.e.m. (*black line* and *vertical black bars*, respectively) across all participants after normalization.

We considered this dataset because of the similar experimental conditions (i.e., displaying black images on a white background on a screen in a dimly lit room) in the presence of completely different tasks compared to our study. Moreover, EEG data in both experiments were acquired with a 256-channel EGI system, MagStim EGI Inc., Eugene, OR, which allowed us to process the EEG time series from [45, 46] as described in the *Main Text*. Specifically, we downsampled the EEG recordings from 1 kHz to 250 Hz (i.e., the sampling rate adopted in our study; MATLAB function: **`decimate`**) and then filtered, reconstructed, and rereferencing a total of 192 EEG recordings as described in Section 2.2. For every picture, we considered the 500-ms-long block of EEG data beginning at the onset time of the picture’s display and treated these blocks as independent trials. Finally, for each participant, we computed the phase-locking values for every pair of channels in the frequency band ℬ = *θ* and built a time-varying graph as described in Section 2.3 along with a common resting-state graph, 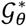. The resultant normalized density of the time-varying graphs and normalized similarity between the resting-state graph and the time-varying graphs are reported in **Fig. S2**.

**Table S1.**
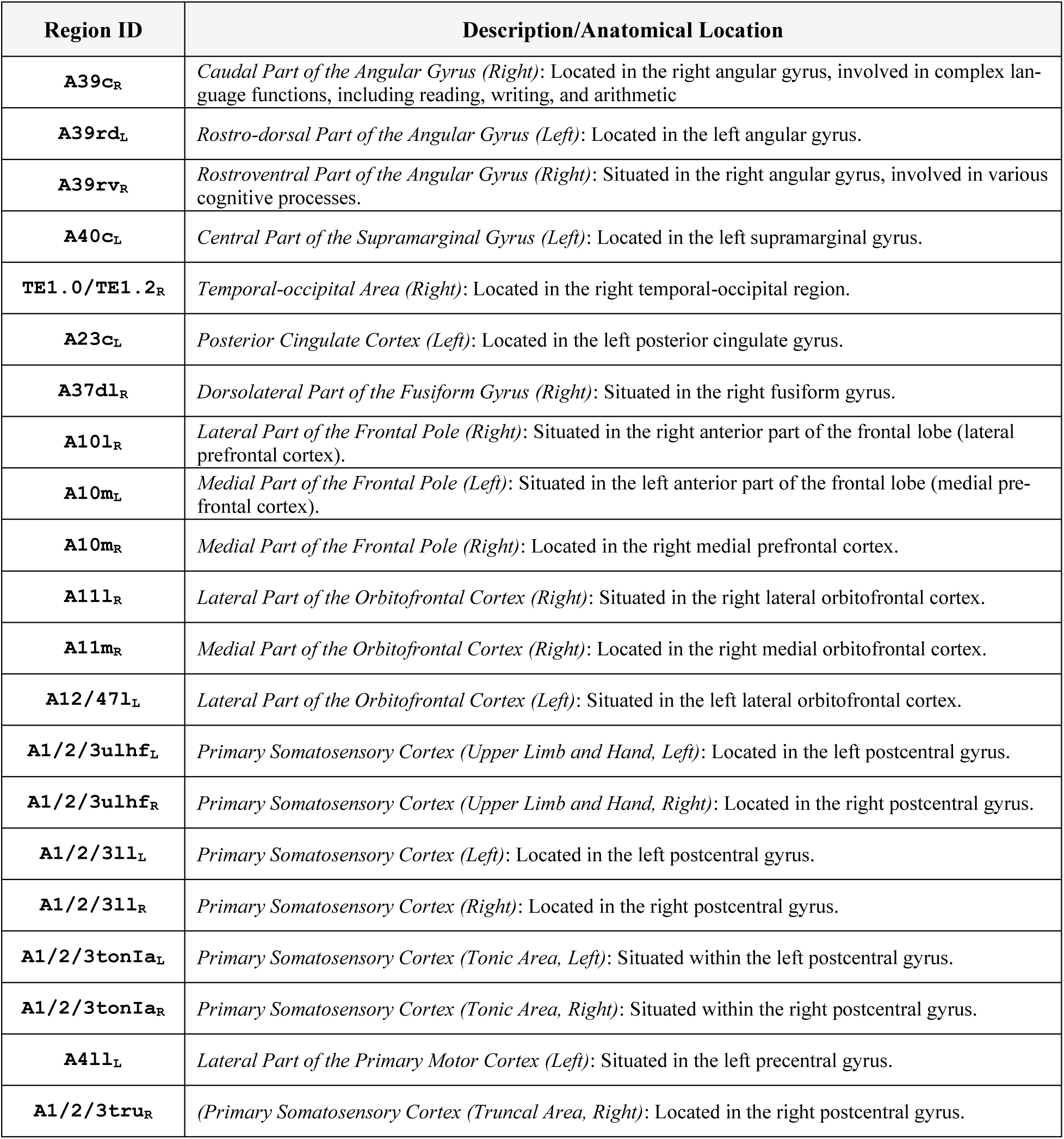
Segmented regions of MNI ICBM 152 atlas [41] with the highest average PageRank centrality calculated across 28 participants during reading. Regions correspond to the labels on the right side of the *red vertical line* in Fig. 4A in *Main Text* and listed from highest to lowest. Average PageRank centrality was calculated for 𝓑 = *θ* across participants and words.

